# Age-associated changes to mouse oocyte meiotic spindle properties revealed through *in situ* measurements

**DOI:** 10.64898/2026.07.01.735913

**Authors:** Marcus A. Begley, Maya Minsky, Karen Schindler

**Affiliations:** Department of Genetics, Rutgers University, Piscataway, NJ 08854; Human Genetics Institute of New Jersey, Piscataway, NJ 08854

## Abstract

Chromosome segregation errors in oocyte meiosis are a leading cause of early miscarriage and congenital disorders in mammals and these errors become more prevalent with advanced maternal age. Although the effects of aging on the functions of critical meiotic proteins and cytoskeletal filaments in oocytes are known, the influence of aging on the force generating capabilities of oocyte spindle components remains largely unexplored. Through the integration of a coarse-grained model and *in situ* experiments, we compare the long-axis mechanical properties of metaphase I (MI) and II (MII) oocyte spindles from reproductively young and old mice. Increased inter-kinetochore distance in aged MII oocytes agree with a model of age-associated cohesion loss, and kinetochore dynamics in these spindles following laser ablation suggest a similar reduction in inter-kinetochore bridge viscosity. Simultaneously, we find that both cohesive and poleward force generators lose stiffness with advanced age in MI spindles. In total, we quantify the extent to which structural spindle components lose their stiffness and viscosity during maternal aging, highlighting the multifaceted impacts of aging on mouse oocyte spindle mechanics.

**Significance Statement:**

- Maternal aging influences mammalian oocyte spindles in numerous ways, yet the impacts of aging on the balance of collective spindle forces remain poorly understood.
- Integrating coarse-grained mechanical modeling with *in situ* measurements of spindle morphology and kinetochore dynamics, we quantify age-associated changes to the viscosities and elastic stiffnesses of oocyte spindle component parts.
- This work provides both a characterization of the effects of aging on force production in mammalian oocyte spindles and a blueprint for future studies of spindle force generation in complex biological contexts.

## Introduction

During eukaryotic cell division, the spindle, a microtubule-based machine, gathers, organizes, and separates a cell’s chromosomes. Successful completion of cell division demands that the spindle generate and apply precise force to each chromosome at the right time. The failure of a cell to properly coordinate these forces can result in daughter cells with the incorrect number of chromosomes, a condition known as aneuploidy. Chromosome segregation is particularly error prone in human oocytes, the female gametes, making aneuploidy a leading cause of birth defects and early miscarriage (Hassold and Hunt, 2001; Nagaoka *et al*., 2012). Importantly, egg aneuploidy and, by extension, female subfertility results from the confluence of mechanically complex factors, such as aging, genetics, and the environment.

Maternal age has long served as a predictor of chromosome segregation fidelity in oocytes, with egg aneuploidy rates in humans rising exponentially after the age of ∼35 years (Hassold and Hunt, 2001; Gruhn *et al*., 2019). This rise correlates strongly with a loss of the cohesin protein complex from chromosomes and an increased stretching of the centromere. It is widely accepted that weakened inter-kinetochore cohesion and more frequent premature sister kinetochore separation result from an age-related diminution of centromeric cohesin (Hodges *et al*., 2005; Chiang *et al*., 2010; Lister *et al*., 2010; Patel *et al*., 2015; Burkhardt *et al*., 2016). Additionally, microtubule dynamics and microtubule-kinetochore attachments are also altered by maternal aging (Shomper *et al*., 2014; Nakagawa and FitzHarris, 2017). Together, these findings imply that poleward forces acting on kinetochores (KTs) change in aging oocytes, alongside changes in cohesive forces that maintain tension between sister and homologous KTs. However, although many of the important ingredients for preserving oocyte quality are known, less is known about how these protein components, combined with proper microtubule dynamics, collectively generate the net force on KTs required to accurately align and segregate chromosomes. Furthermore, how chromosome segregation mechanics in oocytes change with maternal age remains poorly understood. Insights into bulk force generation in metaphase mammalian oocytes could help mechanistically bridge meiotic protein and microtubule kinetics with observed infertility phenotypes.

To investigate how spindle mechanics change with age, we employed a simple mechanical (i.e. coarse-grained) model to estimate changes to the spring constants and drag coefficients of poleward and cohesive spindle forces in MI and MII mouse oocytes during maternal aging. Using this mathematical framework, we track KTs following live-cell kinetochore-fiber (k-fiber) laser ablation, estimate the timescales of k-fiber and inter-KT relaxation, and compute the relative strengths of the viscous and elastic components of both poleward and cohesive forces. We also compare the elastic stiffnesses of the poleward and cohesive forces on KTs by measuring spindle morphology in fixed metaphase oocytes, finding that the balance between poleward and cohesive elastic stiffness changes little with advanced reproductive age. Finally, by integrating the data from these two approaches using our coarse-grained spindle model, we estimate the degree to which advanced maternal age impacts the elastic stiffnesses and viscosities of poleward and cohesive force generation in MI and MII spindles.

Our data indicate that, in both MI and MII spindles, inter-KT viscosity and elastic stiffness decrease by similar amounts with age, indicating a broad increase in the sensitivity of inter-KT linkages to extensile and compressive forces. Furthermore, our data indicate that the elastic stiffness of the poleward pulling force in MI spindles declines with age at a rate comparable to the loss of cohesion. We therefore propose that reduced poleward pulling force and inter-KT viscosity in MI could contribute to age-associated chromosome segregation errors and subfertility.

## Results

### A coarse-grained model for long-axis oocyte meiotic spindle mechanics

To better understand the coordination of chromosome movements in oocyte spindles, we constructed a coarse-grained one-dimensional model of the meiotic spindle (Figure 1, Table 1). The purpose of the model’s simplicity is to enable quantitative estimates of changes to the mechanical properties of mammalian oocyte spindles in wide-ranging and noisy biological contexts such as aging. Specifically, the model provides a readout for the elasticity and viscosity of the forces acting on either side of kinetochores (KTs) during metaphase using data collected from *in situ* assays.

**Figure 1.**
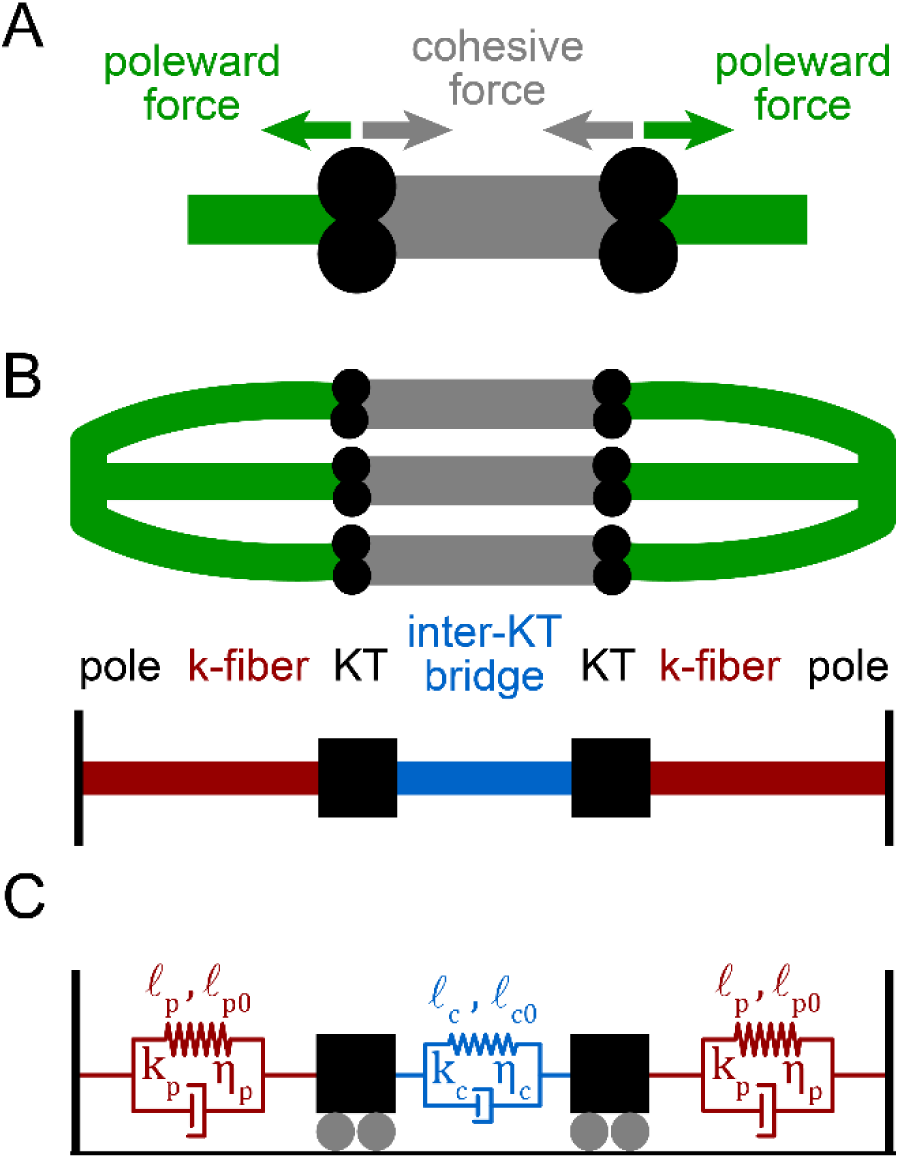
Model of long-axis spindle forces as a series of viscoelastic materials. (A) Schematic illustrating the long-axis forces acting on kinetochores (KTs) (black circles) in a metaphase spindle. (B) Schematic illustrating a spindle coarse-grained into a one-dimensional series of three separate component parts, comprising poleward force generators (red), represented as k-fibers, and a cohesive force generator (blue), joined at kinetochores (black squares). (C) Mechanical spindle model in which the poleward force generators and cohesive force generator are Kelvin-Voigt materials, with elastic (springs) and viscous (dashpots) components, joined at kinetochores (black carts). Variables include the stretched and relaxed lengths of the poleward force generator (*l_p_*, *l_p_*_0_) and the inter-KT bridge (*l_c_*, *l_c_*_0_), as well as the spring constants and drag coefficients of the poleward force generator (*k_p_*, *η_p_*) and inter-KT bridge (*k_c_*, *η_c_*).

**Table 1.**
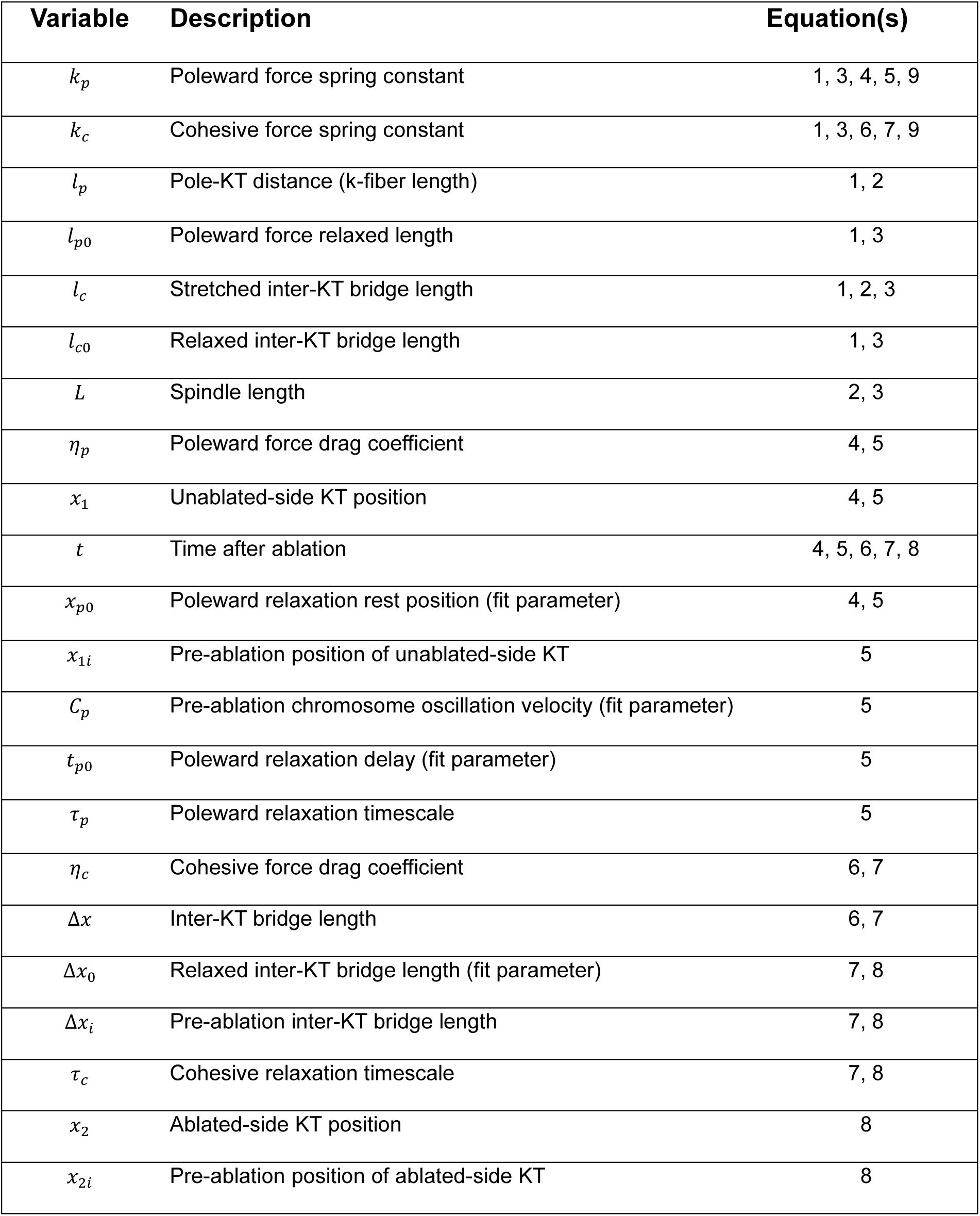
Parameters of the mechanical spindle model.

| Variable | Description | Equation(s) |
| --- | --- | --- |
| $k_p$ | Poleward force spring constant | 1, 3, 4, 5, 9 |
| $k_c$ | Cohesive force spring constant | 1, 3, 6, 7, 9 |
| $l_p$ | Pole-KT distance (k-fiber length) | 1, 2 |
| $l_{p0}$ | Poleward force relaxed length | 1, 3 |
| $l_c$ | Stretched inter-KT bridge length | 1, 2, 3 |
| $l_{c0}$ | Relaxed inter-KT bridge length | 1, 3 |
| $L$ | Spindle length | 2, 3 |
| $\eta_p$ | Poleward force drag coefficient | 4, 5 |
| $x_1$ | Unablated-side KT position | 4, 5 |
| $t$ | Time after ablation | 4, 5, 6, 7, 8 |
| $x_{p0}$ | Poleward relaxation rest position (fit parameter) | 4, 5 |
| $x_{1i}$ | Pre-ablation position of unablated-side KT | 5 |
| $C_p$ | Pre-ablation chromosome oscillation velocity (fit parameter) | 5 |
| $t_{p0}$ | Poleward relaxation delay (fit parameter) | 5 |
| $\tau_p$ | Poleward relaxation timescale | 5 |
| $\eta_c$ | Cohesive force drag coefficient | 6, 7 |
| $\Delta x$ | Inter-KT bridge length | 6, 7 |
| $\Delta x_0$ | Relaxed inter-KT bridge length (fit parameter) | 7, 8 |
| $\Delta x_i$ | Pre-ablation inter-KT bridge length | 7, 8 |
| $\tau_c$ | Cohesive relaxation timescale | 7, 8 |
| $x_2$ | Ablated-side KT position | 8 |
| $x_{2i}$ | Pre-ablation position of ablated-side KT | 8 |

We began developing the model by partitioning the net long-axis force acting on KTs at metaphase into two components: (1) forces acting on KTs from the poleward side (poleward, *p*) and (2) those acting on KTs from the central spindle (cohesive, *c*) (Figure 1A and B). Cohesive forces at MII can be modeled the same as in mitosis, in which sister KT cohesion is often modeled as viscoelastic (Armond *et al*., 2015; Elting *et al*., 2017; Risteski *et al*., 2022). For MI oocytes, both cohesin and chiasmata on chromosome arms provide cohesion between homologs, which provides a restoring force between pairs of sister KTs in a homologous set when the distance between them is stretched. Therefore, like MII, we chose to model the link between homologous pairs of sister KTs in MI spindles as viscoelastic.

Net poleward forces responsible for moving metaphase chromosomes comprise two competing mechanisms: (1) (poleward) forces from the k-fiber and its interactions with nearby microtubules and (2) (anti-poleward) poleward ejection forces (PEFs) (Rieder and Salmon, 1994; Barisic *et al*., 2014; Risteski *et al*., 2021). Metaphase chromosome oscillations in somatic human cells are predominantly driven by PEFs, which can be approximated as scaling linearly with pole-KT distance in the spindle midzone (Joglekar and Hunt, 2002; Ke *et al*., 2009; Armond *et al*., 2015). By assuming a linear dependence of PEFs on pole-KT distance, the total force acting on the poleward side of a KT, a combination of k-fiber pulling force and PEFs, can be approximated as linearly dependent on pole-KT distance, regardless of whether or not the magnitude of k-fiber pulling force is linearly dependent on k-fiber length (Supplemental Figure S1). Recently, Zhu *et al*. (2026) showed that chromosome movements are coordinated in metaphase *Potorous tridactylus* kidney 1 (PtK1) cells, modeling connections between chromosomes as springs that link each chromosome with all others in the metaphase plate. For minor deviations from the metaphase plate, these forces can be approximated as linearly position-dependent, negative poleward elastic forces. Lastly, the spindle responds to small external compressive force applied along its long axis viscoelastically (Itabashi *et al*., 2009), with viscosity dominating its structural response to long-axis forces applied inside the spindle (Shimamoto *et al*., 2011), indicating the importance of a viscous component of the poleward force in our model. Therefore, we modeled the forces acting on KTs from the poleward side as viscoelastic. The elastic component of this poleward force functions as a spring, with a rest length corresponding to the pole-KT distance at which the pulling force of a spindle half on a KT is equal in magnitude to the PEF from that pole.

In total, we modeled long-axis spindle forces as a one-dimensional series of Kelvin-Voigt viscoelastic materials (Osswald and Menges, 2012) with KTs, which we assumed to be point masses, acting as junctions (Figure 1C). This series comprises two k-fibers and their mechanical associations with the larger spindle body that exert poleward force on kinetochores, pulling them apart, separated by an inter-kinetochore (inter-KT) bridge (centromeres in MII spindles) that maintains cohesive force between KTs.

### Morphology and kinetochore elastic force balance are altered in old MII spindles

To study how the balance between poleward and cohesive force elastic stiffnesses might change with advanced maternal age, we compared oocytes from reproductively young and old female mice. The reproductively young mice were 6-12 weeks old, as this is the earliest age at which females are sexually mature. Oocytes from mice ranging from 9-to-11-months of age, have changes to both chromosomal cohesin and MII relaxed inter-KT separation despite having low aneuploidy rates (Koehler *et al*., 2006; Chiang *et al*., 2010; Merriman *et al*., 2012), hinting at potential sub-catastrophic age-related changes to spindle forces in these oocytes. We therefore chose to designate 40-52 weeks of age as reproductively old.

After collecting germinal vesicle-intact, prophase I-arrested oocytes and maturing them *in vitro* to MI, we fixed and immunostained the oocytes and examined spindle morphology using high-resolution confocal microscopy (Figure 2A). In intact bipolar spindles, each sister KT pair in a homologous set is pulled poleward by the spindle/k-fibers and toward the metaphase plate by the inter-KT bridge it shares with the other sister KT pair. In a steady-state metaphase spindle, with static chromosomes aligned at the metaphase plate, the two pairs in the set experience elastic cohesive and poleward forces of equal magnitude:

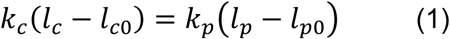

where *k_c_* and *k_p_* are spring constants for the cohesive and poleward forces respectively, *l_c_* and *l_c_*_0_ are stretched and relaxed (as in a monopole) inter-KT distances respectively, *l_p_* is the pole-KT distance, and *l_p_*_0_ is the poleward force relaxed length. Assuming chromosome alignment, pole-KT distance can be approximated as

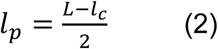

where *L* is spindle length. Substituting Equation 2 into Equation 1 and rearranging gives a ratio for the elastic stiffnesses of the spindle’s cohesive and poleward force generators (*k_c_*⁄*k_p_*):

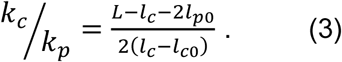

This relationship can be approximated for each class of spindles using measurements from bipolar spindles (*L* and *l_c_*) and monopolar spindles (*l_c_*_0_ and *l_p_*_0_) where both inter-KT bridge tension and opposing pulling from a second spindle pole are minimized (Figure 2).

**Figure 2.**
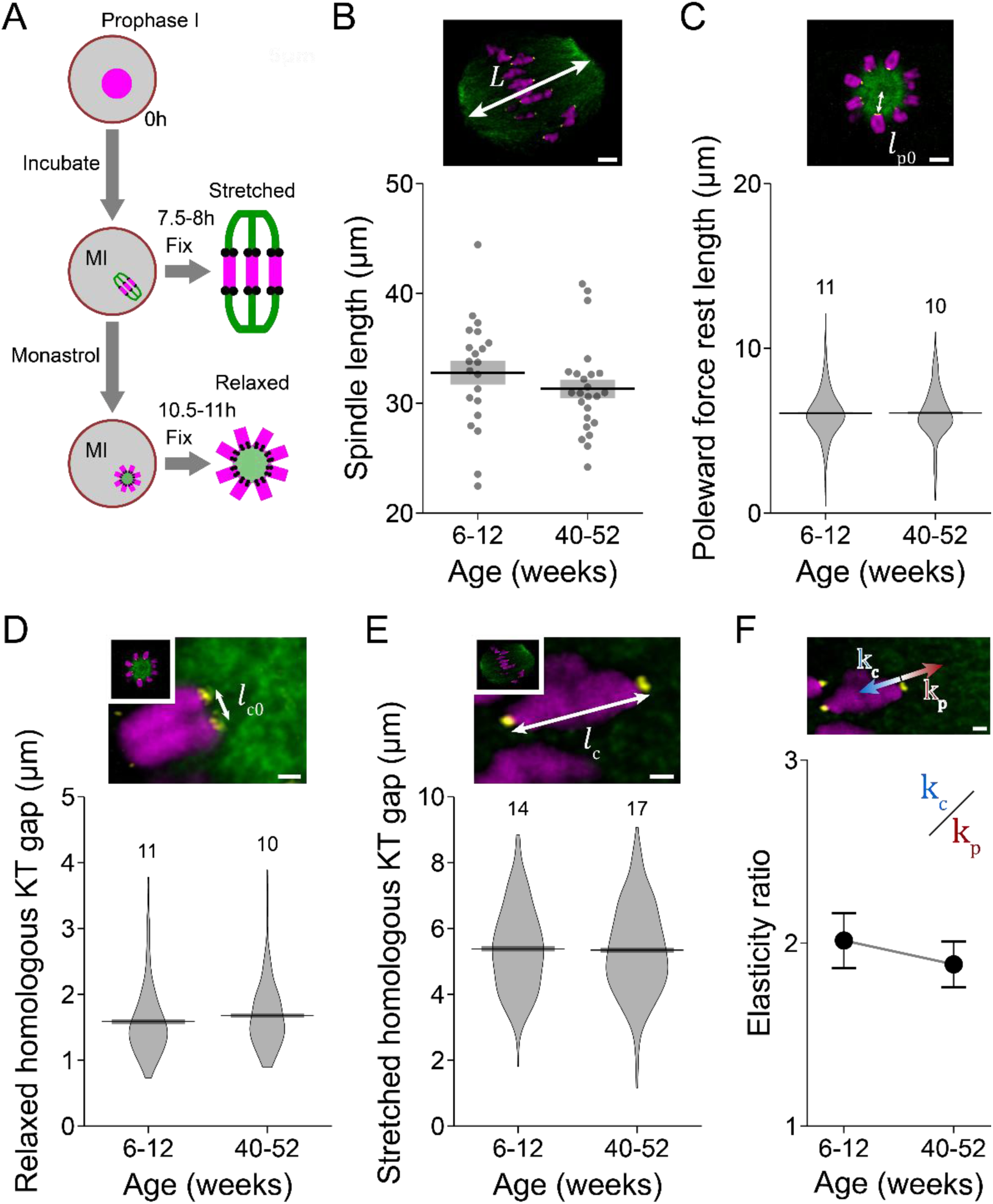
Spindle morphology and kinetochore elastic force balance of MI spindles do not change with age. (A) Schematic describing MI spindle morphology experiments with microtubules in green, kinetochores in black and chromosomes in magenta. (B-E) Spindle morphology data (bottom) collected from young (left) and old (right) mice with example confocal images (top). Spindles were stained to detect tubulin (green), anti-centromeric antigen to detect kinetochores (yellow) and DAPI to stain chromatin (magenta). Measurements include spindle length (B), relaxed pole-KT distance (C), relaxed inter-KT distance (D) and stretched inter-KT distance (E). Insets in (D) and (E) are full images (repurposed from B and C). (F) Relationship between poleward (*k_p_*, red) and cohesive (*k_c_*, blue) spring constants in MI spindles from young and old oocytes. Scale bars are either 5 μm (B-C) or 1 μm (D-F). Horizontal lines (B-E) and dots (F) are mean values and boxes (B-E) and error bars (F) are +/- SEM. Spindle length data (B) was obtained from 12 young mice across 6 experimental replicates and 20 old mice across 4 experimental replicates. All other data (C-E) was obtained from 6 young mice across 3 experimental replicates and 15 old mice across 3 experimental replicates. The number of oocytes used for each measurement are indicated above the data (C-E).

Using this relationship, we found that MI spindle morphology did not change with age (Figure 2, B-E; Table 2), meaning that the *k_c_*⁄*k_p_* ratio also did not significantly change (2.01 ± 0.15 young vs 1.88 ± 0.12 old) (Figure 2F; Table 2). These data suggest that MI spindles maintain a consistent structure and spindle axis KT elastic force balance that are not impacted by maternal age, a surprising result given the relative lack of cohesin and chiasmata on bivalent arms in oocytes from older mice (Hodges *et al*., 2005; Chiang *et al*., 2010; Lister *et al*., 2010; Tachibana-Konwalski *et al*., 2010).

**Table 2.**
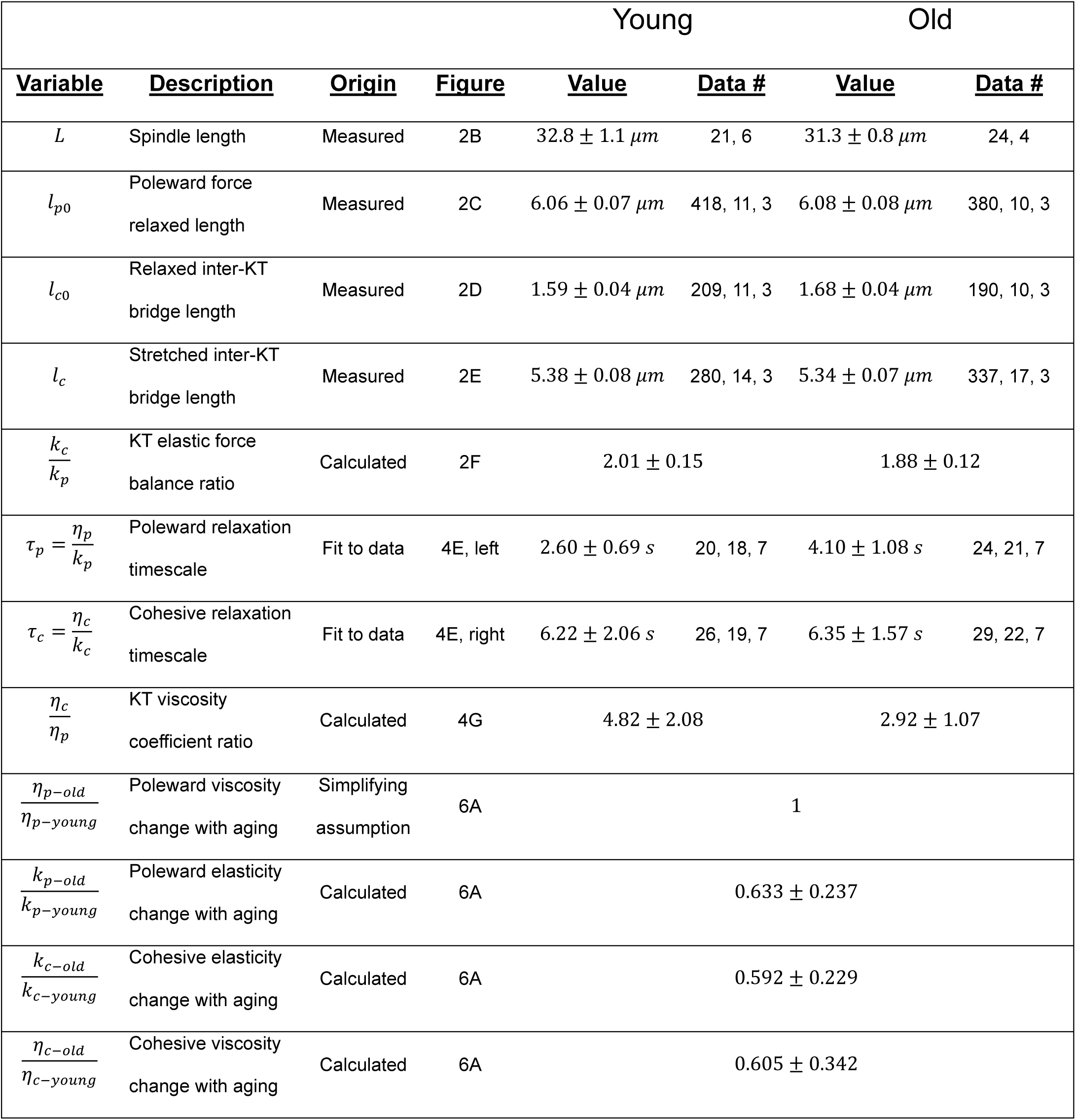
Results of MI spindle morphology measurements and force relation calculations. Values are given as mean +/- SEM and data counts are given as data points, oocytes and experimental replicates.

We next measured the same characteristics of MII oocyte spindles (Figure 3A). Like MI oocytes, we observed no significant change in spindle length (Figure 3B; Table 2) and poleward force relaxed length (Figure 3C; Table 2) with age. Importantly, however, inter-KT bridges were significantly longer in aged oocytes regardless of their tension status (0.90 ± 0.01 *μm* vs. 1.04 ± 0.02 *μm* in monopolar spindles; 1.99 ± 0.02 *μm* vs. 2.19 ± 0.02 *μm* in bipolar spindles) (Figure 3, D and E; Table 2), implying that KT elastic force balance is shifted poleward in these spindles (*k_c_*⁄*k_p_* = 3.52 ± 0.14 vs. 2.48 ± 0.19) (Figure 3F; Table 2). Although it is possible that poleward elastic stiffness also changes with age in MII spindles, we suspect that this poleward shift of KT elastic force balance primarily results from an age-related loss in centromeric stiffness due to decreased centromeric cohesin as previously described (Hodges *et al*., 2005; Chiang *et al*., 2010; Lister *et al*., 2010; Tachibana-Konwalski *et al*., 2010).

**Figure 3.**
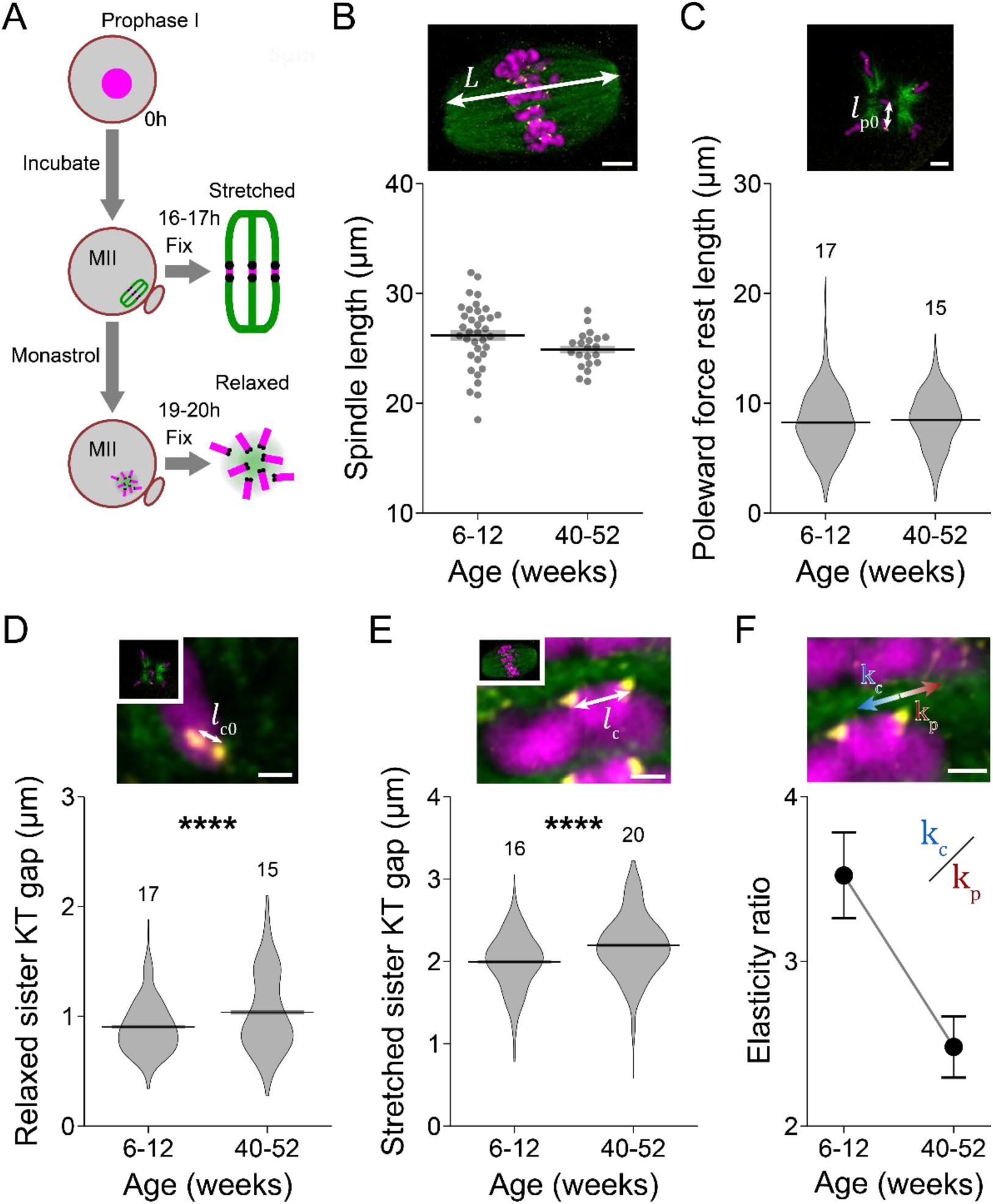
Kinetochore force balance shifts poleward in MII spindles from advanced maternal aged mice. (A) Schematic illustrating MII spindle morphology experiments with microtubules in green, kinetochores in black and chromosomes in magenta. (B-E) Spindle morphology data (bottom) collected from young (left) and old (right) mice with example confocal images (top). Spindles were stained to detect tubulin (green), anti-centromeric antigen to detect kinetochores (yellow) and DAPI to detect chromatin (magenta). Measurements include spindle length (B), relaxed pole-KT distance (C), relaxed inter-KT distance (D) and stretched inter-KT distance (E). Insets in (D) and (E) are full images (repurposed from B and C). (F) Relationship between poleward (*k_p_*, red) and cohesive (*k_c_*, blue) spring constants in MII spindles from young and old oocytes. Scale bars are 5 μm (B-C) or 1 μm (D-F). Horizontal lines (B-E) and dots (F) are mean values and boxes (B-E) and error bars (F) are +/- SEM. Data counts for each measurement are given in Table 3. Spindle length data (B) is obtained from 18 young mice across 9 experimental replicates and 15 old mice across 3 experimental replicates. All other data (C-E) is obtained from 6 young mice across 3 experimental replicates and 15 old mice across 3 experimental replicates. The number of oocytes used for each measurement are given above the data (C-E).

**Table 3.**
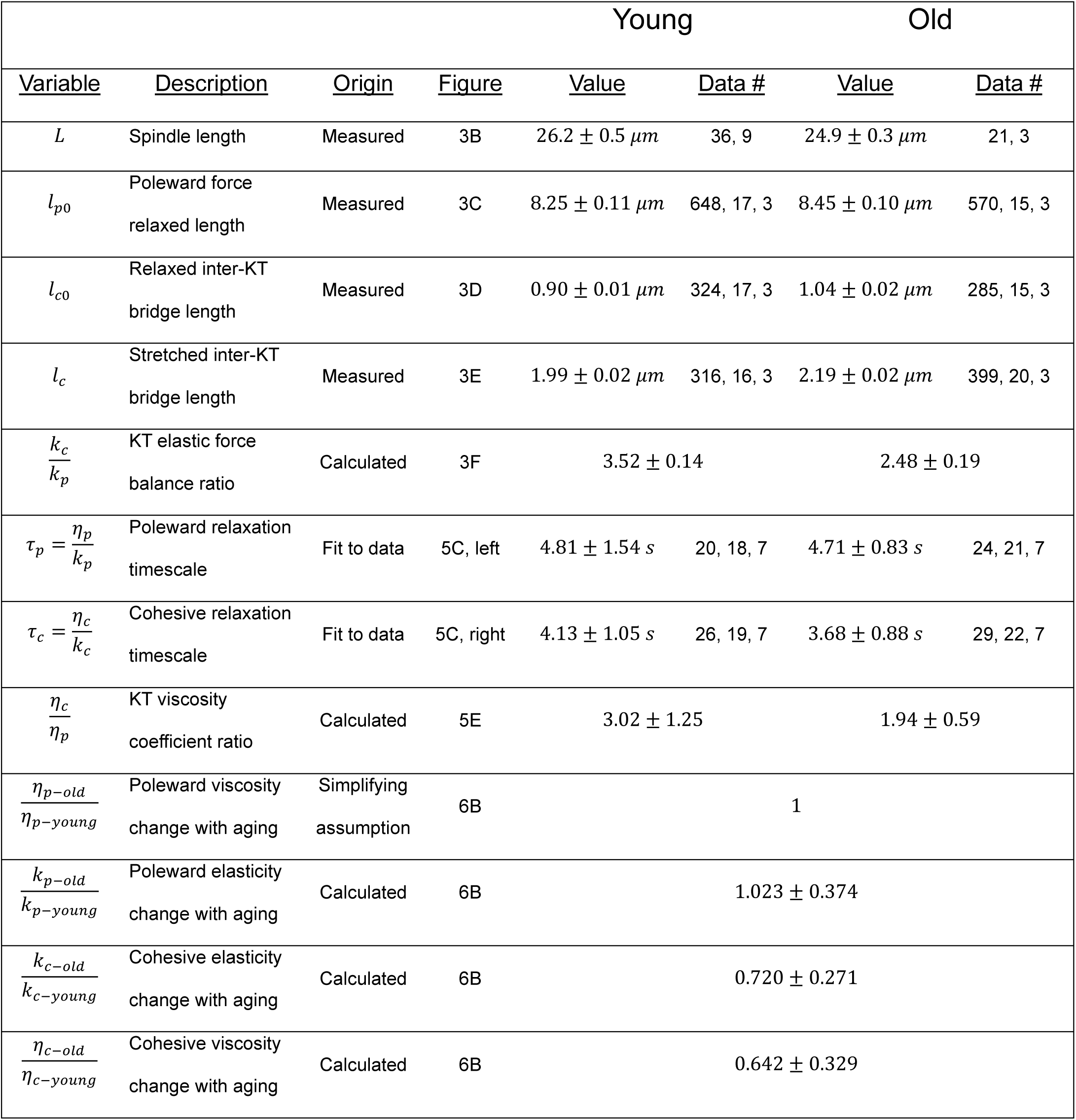
Results of MII spindle morphology measurements and force relation calculations. Values are given as mean +/- SEM and data counts are given as data points, oocytes and experimental replicates.

### Aging affects oocyte poleward and inter-kinetochore viscosities differently

To understand how cohesive and poleward spindle forces each separately change with advanced maternal age, we tracked KTs following k-fiber laser ablation in live MI oocytes (Figure 4, A-D; Supplemental Video SV1). By severing the k-fiber and other nearby microtubules on one side of a KT set, we instantaneously removed the poleward force. As a result, the set/pair moved toward the opposite pole and the inter-kinetochore (inter-KT) bridge relaxed, immediately following ablation (Figure 4, B and C; Supplemental Video SV1). Like previous work in mitotic *Potorous tridactylis* kidney, clone 2 (PtK2) cells, we modeled the inter-KT bridge as a Kelvin-Voigt material and assumed no spindle repair during post-ablation relaxation (Elting *et al*., 2017). However, unlike in PtK2 spindles, in mouse oocyte spindles the unablated k-fiber relaxed on roughly the same timescale as the inter-KT bridge and thus cannot be modeled as fixed (Supplemental Figure S2). Instead, we modeled poleward forces acting on chromosomes as a second Kelvin-Voigt material (Figure 4B). We chose not to consider forces resulting from interactions between the ablated k-fiber stub and spindle environment, because these forces are insignificant to inter-KT bridge relaxation in instances when this relaxation occurred (Elting *et al*., 2017). Assuming all parts of this system are highly viscous, inertia can be ignored and forces are balanced throughout.

**Figure 4.**
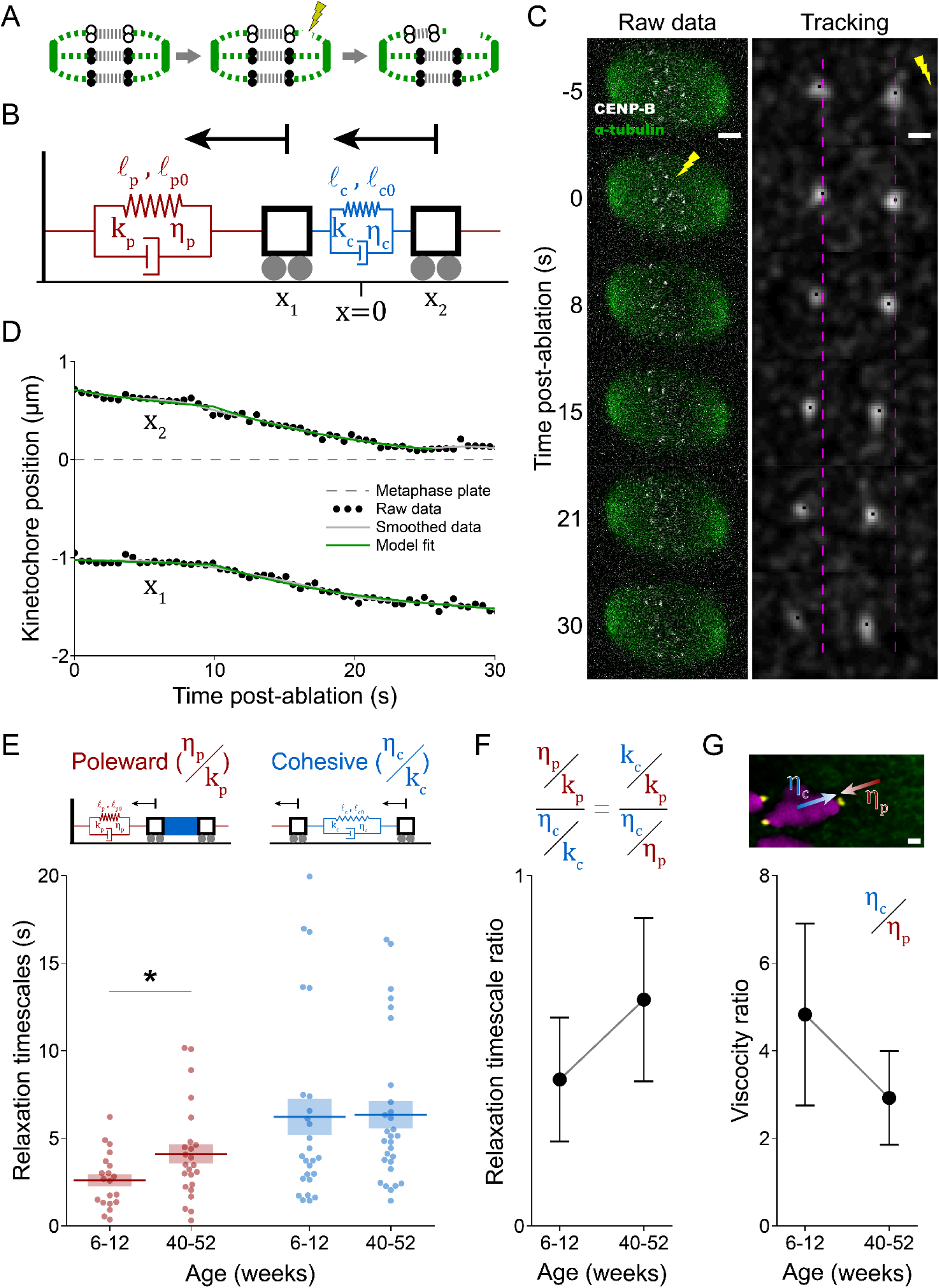
Post-ablation poleward relaxation is gradual in MI spindles from older mice. (A) Schematic showing that k-fiber severance relieves tension in both the inter-KT bridge (gray dashed line) and opposite k-fiber (green dashed line), causing the inter-KT bridge to shorten and both KT pairs (hollow black circles) to move toward the opposite pole relative to the other KTs at the metaphase plate (solid black circles). (B) Model showing that the unablated-side KT pair (*x*_1_, left cart) is pulled toward the unablated-side pole (left wall) by a poleward force generator (red, left) of relaxed length (*l_p_*_0_), stretched length (*l_p_*), spring constant (*k_p_*) and drag coefficient (*η_p_*). Meanwhile, the ablated-side KT pair (*x*_2_, right cart) is pulled toward the unablated-side KT pair by a cohesive force generator (blue, right) of relaxed length (*l_c_*_0_), stretched length (*l_c_*), spring constant (*k_c_*) and drag coefficient (*η_c_*). (C) Example confocal still images of a laser ablation (lightning bolt) experiment in an MI spindle expressing SPY650-tubulin (green) and pIVT-CENP-B-mCherry2 (white) (left) and corresponding Gaussian-blurred tracking video (right). Dashed lines denote pre-ablation KT pair positions (t=0s), and scalebars are 5 μm (left) and 1 μm (right). (D) Post-ablation KT pair position traces (black dots) are smoothed (solid gray lines) and fit to our spindle model (dashed green lines) to estimate relaxation timescales. Data is from the video shown in C and Supplemental Video SV1. (E) Poleward (red, left) and cohesive (blue, right) post-ablation relaxation timescales in young and old oocytes. Each dot corresponds to one video. Data is from 21 young mice across 7 experimental replicates and 35 old mice across 7 experimental replicates. (F) Ratio of mean poleward (*η_p_*⁄*k_p_*) and cohesive (*η_c_*⁄*k_c_*) relaxation timescales. (G) Ratio of inter-KT bridge (*η_c_*) and poleward (*η_p_*) drag coefficients for young and old MI spindles. Example image is repurposed from Figure 2F. Scale bars are 1 μm. Horizontal lines (E) and dots (F-G) are mean values and boxes (E) and error bars (F-G) represent mean +/- SEM. * *p* < 0.05.

First, we examined the unablated k-fiber by tracking the motion of the KT set as a whole, via the position of the unablated-side KT pair (*x*_1_), after k-fiber severance in MI oocytes collected from reproductively young and old females. This set moved poleward as the unablated k-fiber elastically relaxed but was slowed by the local environment’s viscosity (Figure 4, C and D), giving a force balance relationship of

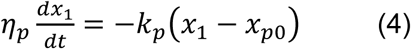

where *η_p_* and *k_p_* are the drag and spring coefficients for poleward force and *x_p_*_0_ is a fit parameter representing the poleward rest position of the unablated-side KT pair. Solving for the position of the unablated-side KT pair (*x*_1_) at time *t* following ablation, gives

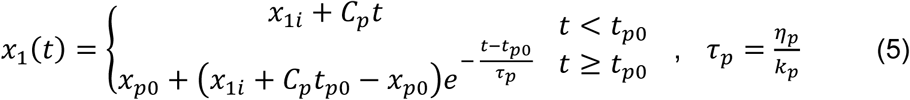

where *x*_1_*_i_* is the position of the unablated-side KT pair at the time of ablation and *τ_p_* is the poleward relaxation timescale. Because we assume the relief of tension of the ablated k-fiber does not instantaneously alter forces on the opposite side of the spindle, we permit unablated k-fiber relaxation to be a piecewise phenomenon where *t_p_*_0_ is the delay in relaxation resulting from the propagation of this mechanical information from one side of the spindle to the other and *x*_1*i*_ + *C_p_t* is the position of the unablated-side KT pair during this delay. *C_p_* and *t_p_*_0_ are arbitrary constants and are fit parameters. Best fits of this form for *x*_1_ (Figure 4D) yielded estimates for the poleward force relaxation timescale (*τ_p_*) in each ablation video (Figure 4E, left side).

We next examined the dynamics of ablated-side KT pairs, which were pulled toward the unablated-side KT pairs as the inter-KT bridge relaxes. This movement was damped by inter-KT viscosity, which we assumed scaled with the relative velocity between the ablated- and unablated-side KT pairs. Altogether, these component forces balanced as

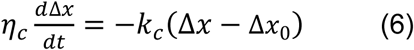

where Δ*x* is the distance between the ablated and unablated KT pairs, Δ*x*_0_ is the inter-KT bridge relaxed length and is a fit parameter, and *η_p_* and *k_p_* are the inter-KT bridge drag coefficient and spring constant, respectively. Solving for Δ*x* gives

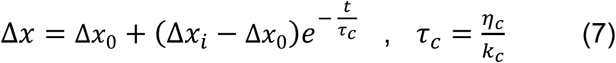

where Δ*x_i_* is the distance between KTs at the time of ablation. Substituting Δ*x* = *x*_2_ − *x*_1_, as well as Equation 5 for *x*_1_, the position of the ablated-side KT pair is

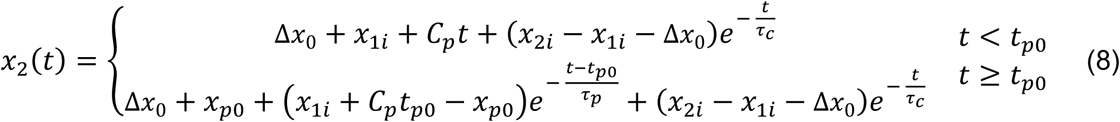

where *x*_2_*_i_* is the position of the ablated-side KT pair at the time of ablation. By computing fits to traces for ablated-side KT pair positions (Figure 4D), we estimated the cohesive force relaxation timescale (*τ_c_*) in each video (Figure 4E, right side).

On average, post-ablation poleward relaxation timescales were slower in spindles from older MI oocytes (2.60 ± 0.69 *s* vs. 4.10 ± 1.08 *s*), possibly due to dysregulated microtubule polymerization kinetics or fewer KT-microtubule attachments (Shomper *et al*., 2014; Nakagawa and FitzHarris, 2017). On the other hand, inter-KT relaxation timing was similar between the two age groups (6.22 ± 2.06 *s* vs. 6.35 ± 1.57 *s*) (Figure 4E). These data were next used, along with the balance between cohesive and poleward elastic stiffness (*η_c_*⁄*η_p_*), to determine the age-dependence of the ratio between cohesive and poleward drag coefficients (*η_c_*⁄*η_p_*) (Eq. 9).

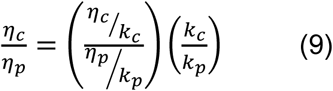

Because poleward and cohesive relaxation dynamics changed unequally with age 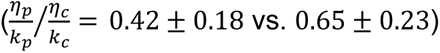 (Figure 4F), but the relative strengths of poleward and cohesive elastic stiffnesses remained constant (Figure 2E), the relative magnitudes of poleward and cohesive viscosities must vary during aging in MI oocytes (*η_c_*⁄*η_p_* = 4.82 ± 2.08 vs. 2.92 ± 1.07) (Figure 4G). Again, this age-dependence could stem from a decrease in cohesive viscosity (*η_c_*), an increase in poleward viscosity (*η_p_*), or a combination of both.

Performing the same experiments in MII oocytes (Figure 5, A and B), we found that neither poleward nor cohesive relaxation dynamics changed greatly with age (Figure 5, C and D). Therefore, because MII KT elastic force balance varies with age (Figure 2J), the relationship between poleward and cohesive viscosities must show a similar dependency on age (*η_c_*⁄*η_p_* = 3.02 ± 1.25 vs. 1.94 ± 0.59) (Figure 5E). In short, k-fiber ablations in MII oocytes revealed centromere viscosity, like centromere stiffness, changes with age.

**Figure 5.**
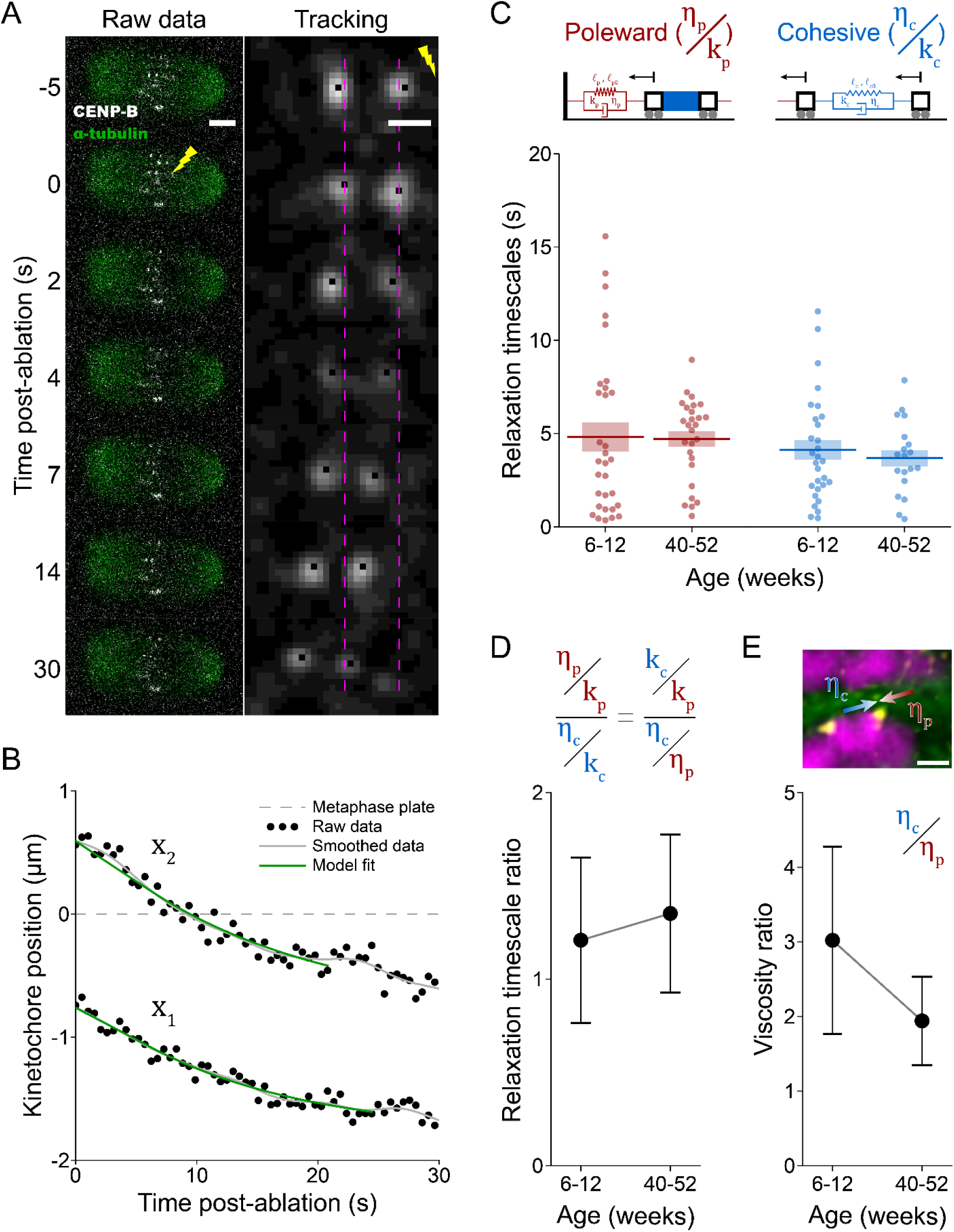
Post-ablation relaxation dynamics in MII spindles do not change with age. (A) Example of confocal images of laser ablation (lightning bolt) in an MII spindle expressing SPY650-tubulin (green) and pIVT-CENP-B-mCherry2 (white) (left) and corresponding Gaussian-blurred tracking video (right). Dashed lines denote pre-ablation KT positions (t=0s), and scalebars are 5 μm (left) and 1 μm (right). (B) Post-ablation KT position traces (black dots) are smoothed (solid gray lines) and fit to our spindle model (dashed green lines) to estimate relaxation timescales. Data is from the video shown in A and Supplemental Video SV2. (C) Poleward (red, left) and cohesive (blue, right) post-ablation relaxation timescales in young and old oocytes. Each dot corresponds to one video. Data is obtained from 23 young mice across 8 experimental replicates and 40 old mice across 8 experimental replicates. (D) Ratio of mean poleward (*η_p_*⁄*k_p_*) and cohesive (*η_c_*⁄*k_c_*) relaxation timescales. (E) Ratio of inter-KT bridge (*η_c_*) and poleward (*η_p_*) drag coefficients for young and old MII spindles. Example image is repurposed from Figure 3F. Scalebars are 1 μm. Horizontal lines (C) and dots (D-E) are mean values and boxes (C) and error bars (D-E) represent mean +/- SEM.

## Discussion

Collectively, our data yield estimates for the ratios of the cohesive and poleward spring constants (*k_c_*⁄*k_p_*), the poleward drag coefficient and spring constants (*η_p_*⁄*k_p_*), and the cohesive drag coefficient and spring constant (*η_c_*⁄*k_c_*) in mouse oocytes. Therefore, with three equations and four unknowns, an empirical solution for how each coarse-grained long-axis physical spindle characteristic changes during aging is unattainable. Examining MI spindles from fixed oocytes highlights this point. That is, in MI spindles, we found that the relative strengths of cohesive and poleward elastic stiffnesses do not change with age (Figure 2F), meaning that inter-KT bridges might be equally stiff in aged MI oocytes as in young oocytes. However, because spindle morphology measurements from fixed oocytes alone cannot separate age-associated changes in cohesive elastic stiffness (*k_c_*) from those of poleward elastic stiffness (*k_p_*), it remains possible that, due to age-associated changes to bivalent structure alongside changes to microtubule dynamics and KT-microtubule attachments (Shomper *et al*., 2014; Nakagawa and FitzHarris, 2017), cohesive and poleward elastic stiffnesses both decrease similarly in MI spindles with age. To address this limitation, we integrated the data presented above with what is known about the effects of aging on oocyte spindles to form a hypothetical model for how these intrinsic properties change during aging.

In *Xenopus* egg extracts, the effective viscosity of the spindle interior is over one hundred times greater than that of the metaphase cytoplasm (Shimamoto *et al*., 2011), meaning that poleward viscosity (*η_p_*) is likely dominated by microtubule crosslinking and density within the spindle, rather than the viscosity of the ooplasm. Although a direct midzone microtubule density measurement has not been made in mouse oocytes, spindles in MI oocytes from young and old mice show no difference in tubulin abundance (Nakagawa and FitzHarris, 2017). Combined with the similarity between old and young MI spindle lengths (Figure 2C), these data suggest that average MI spindle microtubule density remains constant across age groups. Similarly, tubulin density does not significantly decrease in MII oocyte spindles from humans younger than 36 versus older than 39 years of age (Coticchio *et al*., 2013). For these reasons, we chose the simplifying assumption that poleward viscosity (*η_p_*) remains constant with maternal age and estimated the degrees to which each of the remaining metaphase spindle elasticity (*k_c_*, *k_p_*) and viscosity (*η_c_*) parameters change during aging in MI and MII oocytes (Figure 6; Tables 2 and 3), which are discussed below.

**Figure 6.**
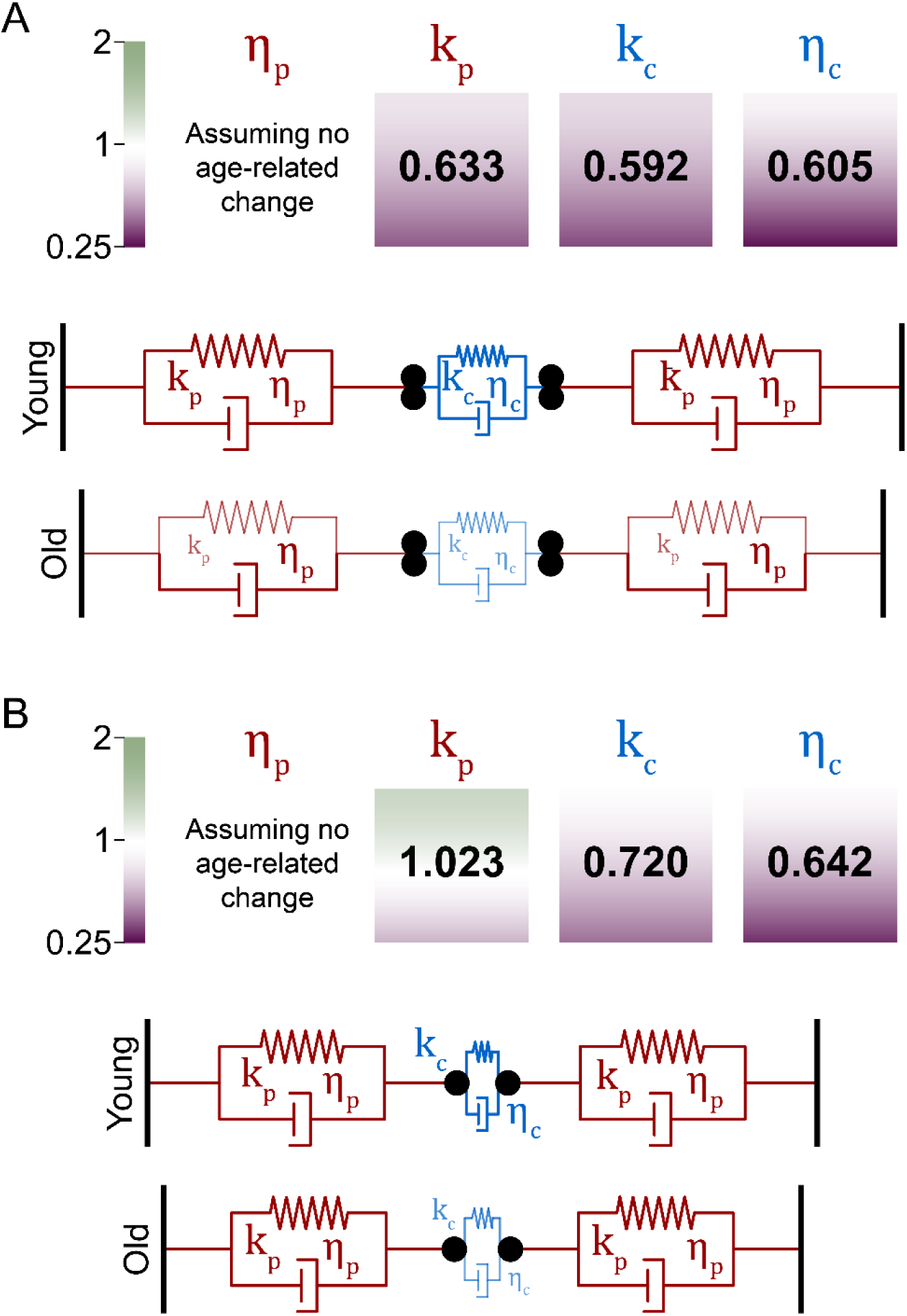
Quantitative estimations of age-associated changes of poleward and inter-KT elasticity and viscosity for MI (A) and MII (B) spindles. (Top row) Values for each coefficient represent the fractional change from young to old (*old*⁄*young*). The opacity of each square is the calculated value +/- (top/bottom) propagated SEM, using a log_4_ scale. (Bottom row) Coarse-grained spindle model depicting age-associated mechanical changes. Spindle geometry is scaled to average spindle length (*L*) and stretched inter-KT distance (*l_c_*) values. Line width, font size, and opacity are scaled by the fractional change from young to old (*old*⁄*young*) of the corresponding spring or drag coefficient.

### Changes in MI spindles

Assuming constant poleward viscosity, the data supports a model that the slower post-ablation poleward movement (*η_p_*⁄*k_p_*) of chromosomes in MI oocytes from reproductively old mice results from a decline in the average poleward force elastic stiffness (*k_p_*) (Figure 4E, left; Table 2). This result both fits the model by which MI chromosome dynamics are influenced by the increased prevalence of KTs lacking microtubule attachments in older MI spindles (Shomper *et al*., 2014; Nakagawa and FitzHarris, 2017), due to dysregulated microtubule dynamics, and provides a rough quantification of this effect (*k_p_*_− *old*_⁄*k_p_*_−*young*_ = 0.63 ± 0.24). Next, because the ratio of cohesive and poleward elastic stiffnesses remained constant in MI spindles (Figure 2F; Table 2), it would logically follow that cohesive elastic stiffness decreases with age (*k_c_*_− *old*_⁄*k_c_*_−*young*_ = 0.59 ± 0.23), consistent with both the loss of cohesin and the reduction and more distal location of chiasmata on homologs observed in aged MI mouse oocytes (Henderson and Edwards, 1968; Hodges *et al*., 2005; Chiang *et al*., 2010; Lister *et al*., 2010; Tachibana-Konwalski *et al*., 2010). Finally, because aging has no apparent effect on post-ablation inter-KT bridge relaxation timescales (*η_c_*⁄*k_c_*) in MI (Figure 4E, right; Table 2), cohesive viscosity must therefore lessen with age comparable to cohesive elastic stiffness (*η_c_*_− *old*_⁄*η_c_*_−*young*_ = 0.60 ± 0.34), suggesting that inter-KT bridges might be more easily deformed from extensile and compressive forces in aged oocytes. Collectively, data obtained in MI oocytes suggests a model where aging leads to a comparable reduction in poleward force elastic stiffness (*k_p_*), inter-KT elastic stiffness (*k_c_*), and inter-KT viscosity (*η_c_*) (Figure 6A; Table 2). Future experiments quantifying KT movements in metaphase I and anaphase I with high temporal resolution would help test whether this apparent age-associated loss of poleward pulling force specific to MI oocytes manifests as altered metaphase chromosome oscillations or slowed chromosome segregation in anaphase I in aged oocytes.

### Changes in MII spindles

We performed a similar analysis to determine the effects of aging on these three parameters in MII oocytes (Figure 6B). Unlike in MI oocytes, we observed no difference in post-ablation poleward relaxation timescales in MII spindles from young and old mice (Figure 5C, left), meaning that, assuming that poleward viscosity does not change with age, poleward elastic stiffness also remains constant during aging (*k_p_*_− *old*_⁄*k_p_*_−*young*_ = 1.02 ± 0.37). This intriguing result hints at either differences in the activities of critical spindle proteins or in the reliance on different spindle-building pathways between MI and MII oocytes. The apparent age-independence of poleward elastic stiffness implies that the observed difference in the poleward and cohesive elastic force balance (*k_c_*⁄*k_p_*) between young and old MII oocytes reflects an age-associated decrease in cohesive elastic stiffness (*k_c_*_− *old*_⁄*k_c_*_−*young*_ = 0.72 ± 0.27) (Figure 3F), likely resulting from the loss of centromeric cohesin in aged oocytes (Hodges *et al*., 2005; Chiang *et al*., 2010; Lister *et al*., 2010; Patel *et al*., 2015). This interpretation is further supported by our observation that the most significant age-associated change to MII spindle morphology is increased inter-KT distance in both monopolar and bipolar spindles (Figure 3, D and E). Although a previous report found no difference in inter-KT distance between bipolar MII spindles from mice of 11-13 weeks and 47-50 weeks of age, those data were collected from live oocytes expressing CENP-C-EGFP to visualize KTs (Kouznetsova *et al*., 2022). Our data closely matches that of prior research showing a significant age-associated change in bipolar MII inter-KT distance, using methods of immunostaining fixed oocytes with anti-centromeric antibody (CREST) similar to ours (Chiang *et al*., 2010). Somewhat surprisingly, post-ablation inter-KT relaxation timescales (*η_c_*⁄*k_c_*) were minimally affected by aging in MII oocytes (Figure 5D). This means that, like MI spindles, cohesive viscosity and elastic stiffness change similarly with age in MII spindles (*η_c_*_− *old*_⁄*η_c_*_−*young*_ = 0.64 ± 0.33), suggesting that centromeres become more susceptible to extensile and compressive deformations with advanced age. In total, MII oocyte data suggests a model in which poleward force (*k_p_*, *η_p_*) changes minimally for MII spindles, but inter-KT bridge stiffness (*k_c_*) and viscosity (*η_c_*) both significantly decrease with advanced age (Figure 6B; Table 3).

### Mechanical differences between aged MI and MII spindles

Interestingly, assuming poleward spindle viscosity does not change with age, poleward pulling force on KTs changes differently with advanced age between MI and MII spindles hinting at possible differences in microtubule nucleation and organization between these two spindle types. Lending support for this hypothesis is the different protein requirements for MI and MII mouse oocyte spindle construction. For example, HSET is a minus-end-directed kinesin that collects and organizes microtubule minus-ends into asters *in vitro* (Henkin *et al*., 2022). It thereby regulates the timing and rate of spindle bipolarization, as well as the size of spindle poles in MI mouse oocytes (Mountain *et al*., 1999; Bennabi *et al*., 2018). However these MI morphological alterations are minor compared to the loss of spindle bipolarity and minus-end organization found in MII oocytes when HSET activity was blocked by injection of anti-HSET antibodies (Mountain *et al*., 1999). These data suggest that minus-end organization may be a more critical function in MII spindles.

Differences between human and mouse oocytes further support this hypothesis. Interestingly, human oocytes, which lack acentrosomal microtubule organizing centers (aMTOCs) and therefore must actively sort microtubules nucleated elsewhere in the spindle, are deficient in HSET expression (So *et al*., 2022). In this case, pole-focusing is provided by the NuMA/dynein/dynactin complex, demonstrated by the inability of human oocytes to form MI spindles when NuMA and HAUS6, a component of the augmin complex, are simultaneously knocked down via RNA interference (So *et al*., 2022). This co-depletion represents the concurrent impairment of two spindle-building pathways, as augmin is required to form microtubule networks near chromosomes in spindles from acentrosomal *Xenopus* egg extracts (Gouveia *et al*., 2023). Indeed, *HAUS6* mutations have been identified in infertile patients who have disrupted MI spindle bipolarity and pole organization (Wu *et al*., 2024). In this context, the apparent differences in age dependence in poleward pulling force elasticity between MI and MII spindles may stem from differences in the rates at which microtubules are seeded from aMTOCs versus from chromatin, even though both MI and MII oocytes contain aMTOCs in mouse. It would be interesting to compare the poleward pulling force, or lack thereof, in MI and MII mouse oocytes deplete of pericentrin, a critical aMTOC protein (So *et al*., 2022), to test the degree to which aMTOCs are required for long-axis force generation in MI and MII spindles.

Altogether, our data demonstrates that long-axis spindle forces are sensitive to maternal age for both MI and MII mouse oocytes. However, while we have presented and employed a method for quantitatively estimating changes to long-axis spindle forces in response to complex pressures, such as aging, we note some limitations. First, forces acting on chromosomes in metaphase vary depending on numerous factors not included by our model, such as the sizes of the chromosomes, their positions in the spindle, and local microtubule orientations (Kelleher *et al*., 2024; Takenouchi *et al*., 2024; Mishina *et al*., 2025). Here, we chose to focus on changes to the most coarse-grained of long-axis spindle properties as a starting point. Nevertheless, these factors constitute a likely source of variability and noise in our data. Another constraint of this study is that evaluating the age dependence of each spring constant and drag coefficient requires simplifying assumptions regarding the age dependence of one of these quantities. Based on previous research, it seems likely that poleward spindle viscosity changes little with age in oocytes (Coticchio *et al*., 2013; Nakagawa and FitzHarris, 2017), but this is not easily measured *in situ*. Relatedly and importantly, the data presented here is meant to compare these spindle properties between oocytes from differently aged mice rather than represent absolute measured values for these parameters. *In vitro* experiments that more directly measure inter-KT elasticity and/or viscosity, similar in nature to previous measurements of oocyte chromosome stiffness (Hornick *et al*., 2015; Liu *et al*., 2025), would help reduce the demand for simplifying assumptions and add further clarity to the age dependence of these bulk mammalian oocyte spindle properties.

In summary, our data implies that links between pairs of sister KTs in MI spindles and centromeres themselves in MII spindles become both less stiff and less viscous with advanced maternal age. We therefore posit that aging renders inter-KT bridges in mouse oocyte spindles more susceptible to deformation from both extensile and compressive meiotic forces. Furthermore, poleward pulling force appears to be reduced in MI oocyte spindles from mice of advanced maternal age, potentially impairing chromosome alignment and segregation. More broadly, we speculate that our coarse-grained model and assays can be generally applied to quantify spindle forces in a wide range of noisy cellular systems.

## Materials and Methods

### Oocyte collection

All experiments used C57BL/6J mice (Jackson Lab #000664) and adhered to Rutgers University Institutional Animal Use and Care Committee policies and guidelines (Protocol #201702497). All mice were housed in the same room which maintains a 12-hour light/dark cycle, as well as constant temperature and humidity. Food and water were provided *ad libitum*. Mice were injected with 5 I.U. pregnant mare serum gonadotropin (PMSG) (Lee BioSolutions #493-10) 2 days prior to the collection of oocytes arrested in prophase I (Blengini and Schindler, 2018). Oocytes were collected in minimum essential media (MEM) (Sigma-Aldrich #M0268) containing 2.5 μm milrinone (Sigma-Aldrich #M4659), which prevents meiotic resumption, and were subsequently transferred to Chatot, Ziomek, and Bavister (CZB) media (Chatot *et al*., 1989) with milrinone to hold the arrest prior to microinjection.

### Oocyte maturation, fixation, immunofluorescence and imaging

Oocytes were matured in CZB media without milrinone for 7.5-8.5 h for MI and 16-17.5 h for MII. For monopolar spindle induction, oocytes were cultured an additional 3 h in CZB containing 100 μM Monastrol (MilliporeSigma #M8515), dissolved in dimethyl sulfoxide (DMSO) (MilliporeSigma #472301), in center-well organ culture dishes. Subsequently, oocytes were fixed in phosphate-buffered saline (PBS) containing 2% paraformaldehyde (PFA) (Sigma-Aldrich #P6148) and stained with a Alexa 488-conjugated tubulin antibody (1:100; Cell Signaling Technology #5063S) and anti-centromeric antibody (CREST) (1:30; Antibodies Inc. #15-234) to visualize kinetochores, before being mounted to slides in 10 μL of VectaShield (Vector Laboratories #H-1000) containing 4’,6-Diamidino-2-Phenylindole, Dihydrochloride (DAPI) (1:170; Life Technologies #D1306).

Slides were imaged using a 63X 1.40 NA oil immersion objective (Leica) mounted to a Leica TCS SP8 confocal fluorescence microscope with a Leica DMi8 stand. Excitation lasers with peaks at 405 nm (DAPI), 488 nm (tubulin) and 552 nm (CREST) were all paired with HyD photodetectors. Images were taken using Leica Application Suite X (LASX) software. Each image was captured as a stack of optical z-sections using a 290-300 nm z-step size, 95.55 μm pinhole diameter, 3-5 zoom factor, 1024-2048 p resolution, and 200-400 Hz acquisition speed. Stacks were collected in Lightning mode and the deconvoluted images were used for analysis.

### Microinjection, live imaging and laser ablation

For live kinetochore visualization, oocytes were microinjected with 570 ng/μl mRNA coding for pIVT-CENP-B-mCherry2 using a Xenoworks microinjector (Sutter Instruments) as previously described (Stein and Schindler, 2011). To maintain prophase I arrest, oocytes were suspended in MEM containing milrinone during microinjection and in CZB supplemented with milrinone for at least 3 h following microinjection. Injected oocytes were then released from cell cycle arrest by transfer into CZB without milrinone in a glass-bottom dish (Ibidi #81817). To label spindles, CZB included SPY650-tubulin dye (1:1000) (Cytoskeleton Inc. #CY-SC503).

Live imaging was performed using a 63X 1.40 NA oil immersion objective (Leica) mounted to a Leica TCS SP8 confocal fluorescence microscope with a Leica DMi8 stand, with samples maintaining 37°C and 5% CO_2_ within a humidified TokaiHit STX stage-top incubator. Excitation lasers with peaks at 552 nm (CENP-B) and 638 nm (tubulin) were both paired with HyD photodetectors. For fixed oocyte imaging, video acquisition was managed by LASX. Videos were taken using a 2.5 zoom factor, 95.55 μm pinhole diameter, 1000 Hz acquisition speed, 512 p resolution, and 520 ms frame rate. In-video laser ablation was performed via a galvo-steered Andor MicroPoint Laser Illumination and Ablation System (Oxford Instruments), delivering 30 3-ns pulses of 425 nm light at 16 Hz. Ablations were conducted through Andor iQ3 software in coordination with LASX-managed video acquisition.

### Fixed spindle measurements

Kinetochore positions and spindle length were measured from images using Imaris (Oxford Instruments). Spindle pole positions were identified by eye, and the line tool was used to measure spindle length. The Spots tool was used to mark kinetochores and output their positions. These positions were organized within the Spots tool, with each set of homologous sets of kinetochores (for MI) or pairs of sister kinetochores (for MII) separated into individual classes. Inter-kinetochore distances (*l_c_*_0_ and *l_c_*) and poleward force rest lengths (*l_p_*_0_) were then calculated from kinetochore positions using a home-written Python script. Taking advantage of monopolar spindle geometry, poleward force rest length was defined as the distance between an individual KT (MII) or sister pair (MI) and the average KT position for that image.

### Video processing and kinetochore tracking

To streamline analysis, live-oocyte videos were edited using Fiji. Videos were cropped, brightness and contrast adjusted linearly, and files converted to TIFF stacks (Supplemental Videos SV1-SV2). All subsequent video processing and analysis was conducted using a home-written semi-automated object tracking Python program. For a specific video, this program begins by applying a customizable Gaussian blur to the video using binomial coefficients (Marchand and Marmet, 1983; Aubury and Luk, 1996). Next, a pop-up window displaying the blurred image of the video’s first frame appears and the user marks both kinetochores, before the program automatically tracks these two objects through the video. Once tracking was complete, the program outputs a CSV file of kinetochore positions, and a TIFF stack of the blurred tracking video, with kinetochores labeled in each frame by single black pixels (Figure 3C; Supplemental Videos SV1-SV2, bottom left corner).

### Post-ablation relaxation analysis

All post-ablation KT dynamics analysis was conducted in a home-written Python program. First, a video’s KT position traces were smoothed using a five-point binomial filter (Marchand and Marmet, 1983; Aubury and Luk, 1996), and all subsequent steps used these smoothed positions. Then, positions were averaged from the first 10 frames in the video for each KT (MII) or pair of sisters (MI) to estimate their initial positions and a line was drawn between them as an estimation of the spindle’s long axis. All future KT positions were then projected onto this line, since we were only concerned about long-axis force components. Additionally, position values were then shifted, so that *x* = 0 (representing the metaphase plate) was the midpoint between the two initial KT positions. Pre-ablation KT positions were defined as the KT positions from the frame immediately prior to the initiation of ablation. Finally, best fits of the forms given by Equation 5 and Equation 8 were calculated for post-ablation KT positions (Figure 4D and 5B, yielding values of poleward (*τ_p_*) and cohesive (*τ_c_*) relaxation timescales. For each data set (e.g. poleward relaxation timescales in young MI oocytes), obviously incorrect values (e.g. 1000 min) were removed, before filtering out statistical outliers using a two-sided Grubbs test with 95% confidence. Data omitting statistical outliers was presented (Figure 3E, 3H) and then used for subsequent calculations in Figures 3 and 4.

### Statistical analysis

Statistical significance was determined by unpaired two-sided Student’s t-tests, performed in Python, with *p* < 0.05 considered statistically significant. All error on measurements is presented as mean +/- SEM. For metrics combining experimental data from multiple measurements (Figure 2F, 3F, 4F-G, 5D-E, 6), error is propagated using Equation 10 for addition/subtraction and Equation 11 for multiplication/division,

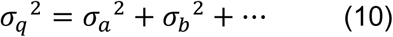

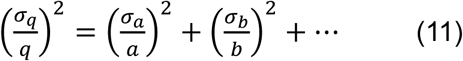

where *σ_q_* is the propagated standard error for calculated value *q*, *σ_a_* is the standard error on mean measured value *a*, *σ_b_* is the standard error on mean measured value *b*.

## Abbreviations

MI: Metaphase I
MII: Metaphase II
KT: Kinetochore
K-fiber: Kinetochore-fiber
PEFs: Poleward ejection forces
aMTOCs: Acentrosomal microtubule organizing centers

## Acknowledgements

We thank members of the Schindler Lab for their helpful comments and suggestions, and Dr. Mary Elting (NCSU) for feedback on the project and manuscript. The pIVT-CENP-B-mCherry2 construct is a gift from Dr. Michael Lampson (U Penn). This work was supported by NIH grant 2R35GM136340 to K.S. and M.A.B. is funded by the NIH IRACDA program (5K12GM093854).

## Author contributions

M.A.B and K.S. designed the experiments; M.A.B performed the experiments; M.A.B. and M.M. analyzed data; M.A.B. and K.S. drafted the manuscript; All authors reviewed and approved the final manuscript.

## Data and code availability

No proteomics or sequence data was generated as a part of this study. Imaging data and detailed experimental protocols will be made available by the authors upon request. Image processing and data analysis scripts are available via the KarenSchindlerLab GitHub.

