## Supplemental Figure S1 for "Age-associated changes to mouse oocyte meiotic spindle properties revealed through *in situ* measurements"

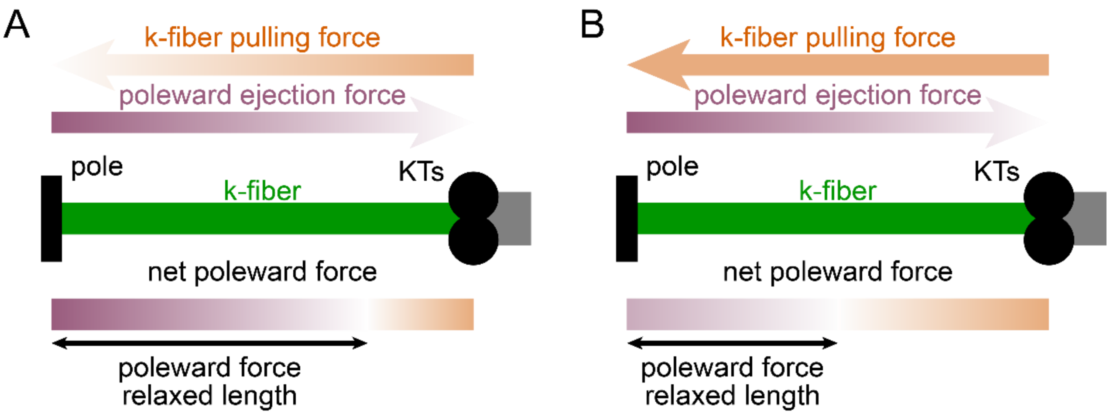


**Supplemental Figure S1.** Both position-dependent (A) and position-independent k-fiber pulling force (B) result in net poleward force on KTs that scale linearly with KT-pole distance, but with different rest positions. Orange and purple represent outward pushing and poleward pulling forces respectively. The spindle pole is represented by as a fixed wall on the left, KTs as black circles and the k-fiber connecting the pole and KTs as a green rod.
