## Supplemental Figure S2 for "Age-associated changes to mouse oocyte meiotic spindle properties revealed through *in situ* measurements"

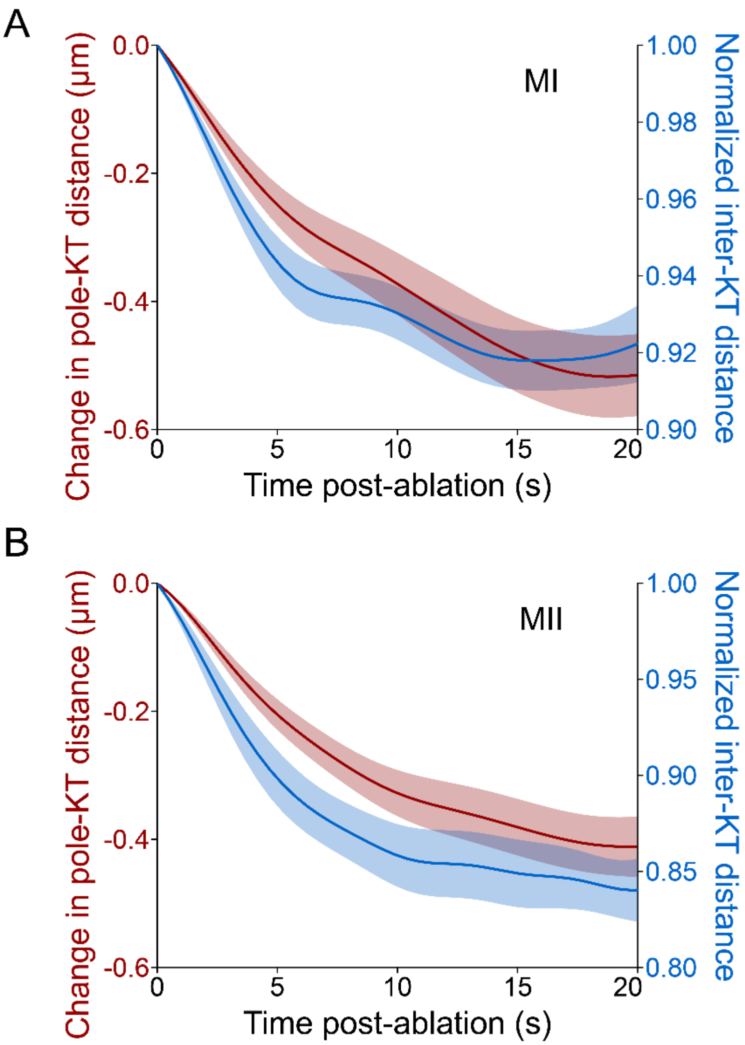


**Supplemental Figure S2.** K-fiber ablation induces poleward (red) and cohesive (blue) elastic relaxation in MI (A) and MII (B) spindles. Poleward relaxation is measured in the post-ablation change in KT-pole distance on the unablated spindle half (left axis) and cohesive relaxation is measured in the post-ablation change in inter-KT distance for the ablated pair (right axis). Data from young mice (6-12 weeks). Shaded regions are mean +/- SEM.
