## Supplemental Videos Legend for "Age-associated changes to mouse oocyte meiotic spindle properties revealed through *in situ* measurements"

**Supplemental Videos SV1 and SV2.** Examples of post-ablation relaxation in an MI (SV1) and MII (SV2) oocyte spindle from a young mouse expressing SPY650-tubulin (green) and pIVT-CENP-B-mCherry2 (white). Times are given as min:sec and scale bars are 5 μm. Red dots immediately pre-ablation label the position of irradiation. Corresponding tracking videos are inset in the bottom left corner with 1 μm scalebars and black pixels denoting the pixels closest to KT positions in each frame.
